# Carrier-Limited Release Risk in Electrospun Nanofibers for Regenerative Endodontics: Constraint Discovery from 25 Formulations

**DOI:** 10.64898/2026.10.29.762783

**Authors:** Nura Brimo, Dilek Çökeliler Serdaroğlu

**Affiliations:** Department of Biomedical Engineering, Duke University, Durham, NC 27708, USA; Department of Biomedical Engineering, Başkent University, Ankara 06790, Türkiye

## Abstract

Regenerative endodontics needs an intracanal carrier that keeps disinfecting across a dressing interval that clinicians schedule at one to four weeks, without harming the stem cells that rebuild the pulp. We ask what one completed 25-formulation electro-spinning campaign of poly(vinylpyrrolidone) (PVP) fibers carrying metronidazole, ciprofloxacin, minocycline and Ca(OH)_2_ can decide, and use machine learning for constraint discovery and decision support rather than as a new predictor. After an audit against the source thesis, logistic regression predicts fine-fiber formation at 96% leave-one-out accuracy, but a one-threshold rule does as well, the gain over the 80% majority baseline is not significant (exact McNemar *p* = 0.125), and accuracy falls to 84% when mixture loading is encoded per component. Diameter models interpolate within formulation families (ridge MAE 193 nm, 95% CI 96–317) and fail across them (*R*^2^ *<* 0). Because the campaign never measured release, we assemble a same-drug, cross-carrier evidence table: 17 bounded metronidazole and ciprofloxacin observations from seven studies, with two-sided censoring. Water-soluble carriers (PVP, PVA) completed release within 80 h (PVP within 8 h); no water-insoluble monolithic carrier reached 90% within its observation window, releasing 1.5–50%; both within-study comparisons separate at matched loading; and blend, core–shell and thick-mat architectures released over 3–28 days. The PVP mats therefore carry a carrier-limited release risk, not a proven ceiling, and one dissolution experiment can confirm or overturn it. A censoring-aware screen keeps three necessarily and one possibly nondominated formulation, and fixed-budget replays show that uncertainty sampling finds failed formulations sooner, except the one it is confidently wrong about.

## 1 Introduction

A tooth whose pulp dies before the root finishes forming cannot be saved by a conventional root-canal filling: the root stays thin, short and prone to fracture. Regenerative endodontic treatment instead disinfects the canal and lets the patient’s own apical stem cells rebuild pulp-like tissue and continue root development [Diogenes and Ruparel, 2017, Albuquerque et al., 2014]. Everything turns on the disinfection step. Clinical guidance schedules the second appointment one to four weeks after the first, with calcium hydroxide or a low-concentration triple antibiotic paste in the canal between visits [American Association of Endodontists, 2022]. Triple antibiotic paste (metronidazole, ciprofloxacin, minocycline) [Hoshino et al., 1996] is toxic to stem cells of the apical papilla at clinical concentrations [Ruparel et al., 2012] and minocycline stains the crown [Kim et al., 2010]. Electrospun fiber mats that carry the same antibiotics at lower doses are a leading alternative [Bottino et al., 2013, Pankajakshan et al., 2016], and the trade-offs between pastes and fibers have been reviewed [Brimo et al., 2022].

Designing such a mat means choosing a polymer, a drug set, a loading and a process, then spinning and assaying the result. A campaign of twenty or thirty formulations is substantial bench work, and machine learning in this area is mostly used as an interpolator inside one material system [Fatahian and Erfani, 2025, Pervez et al., 2023, Woodring et al., 2025, Rostami et al., 2025]. We ask a different question: given one completed endodontic campaign, what can it *decide*? Our answer is that its most useful output is not a prediction inside the design space but a constraint on the design space, stated with the evidence and the uncertainty that a formulation scientist needs before committing the next round of experiments.

### Contributions

1. **EndoDesign-25, audited against the thesis** (§3): 25 outcome-linked formulations with page-level provenance, a twentieth diameter recovered from the thesis, the recipe label separated from physical drug loading, failed runs retained, and every unresolved inconsistency recorded rather than smoothed.
2. **Morphology models with the baselines and uncertainty a 25-row dataset needs** (§4). A one-threshold rule matches the 96% classifier, the accuracy depends on how mixture loading is encoded, and diameter models interpolate within formulation families but fail across them.
3. **A same-drug, cross-carrier evidence table** for metronidazole and ciprofloxacin fibers with two-sided censoring (§5). It removes the drug-identity confound, frames the result as a falsifiable release risk rather than a hard ceiling, and identifies fiber architecture as the lever.
4. **Decision support** (§6–8): a censoring-aware partial order with MIC-error robustness, fixed-budget acquisition replays with an observable stopping rule, and a decision table that maps each outcome of the one missing experiment to an action.

## 2 Related work

### Machine learning for electrospinning and release

Most models map process parameters to fiber diameter within one polymer system, for example small neural networks and response-surface hybrids on designed experiments [Fatahian and Erfani, 2025, Pervez et al., 2023]. Release models are usually fitted within one scaffold family, such as Gaussian-process regression over acetalated-dextran fibers [Woodring et al., 2025] or tree ensembles for doxorubicin-loaded fibers [Rostami et al., 2025]. Our morphology stage differs in its input (composition at fixed process settings), and our release stage asks whether anything transfers to a material system with no release data at all, in the spirit of cross-task benchmarking in materials ML [Dunn et al., 2020]. Machine learning has also been reviewed as a tool for characterizing biomaterials and their mechanical properties [Brimo et al., 2024].

### Endodontic fiber scaffolds

PVP fibers loaded with Ca(OH)_2_ have been used to coat gutta-percha cones as an intracanal carrier and compared with Ca(OH)_2_ paste [Brimo et al., 2026]. Polydioxanone scaffolds carrying metronidazole and ciprofloxacin have been characterized mainly through the antimicrobial and cellular activity of their eluates; at least one study states that the released amount was not quantified [Palasuk et al., 2014]. Quantified release for endodontic fiber mats is therefore scarce, which is why we borrow same-drug evidence from wound, periodontal, ocular and oral-delivery studies (§5).

### Self-driving laboratories and small data

Autonomous platforms pair surrogates with acquisition functions to reduce experiment counts [MacLeod et al., 2020, Shields et al., 2021, Lookman et al., 2019, Settles, 2009]. Retrospective replays can evaluate acquisition policies, but hidden labels must stay hidden from both the acquisition and the stopping decision. Cross-validation on small samples also carries wide error bars [Varoquaux, 2018], so we report intervals, permutation tests and per-family errors rather than one headline accuracy.

## 3 EndoDesign-25: data and provenance

### Sources

Morphology comes from the MSc thesis behind the campaign [Brimo, 2020]; biological endpoints come from the published article [Brimo et al., 2023]. The source study states that it “did not consider the kinetics of drug release”, so no release curve exists for any mat. Fibers were spun from 8% PVP in ethanol at a nominal 0.9 mL h^*−*1^ and 15 cm working distance; composition is the design variable.

### Formulations and outcomes

Single-drug series of metronidazole (MET: 2.5, 5, 7.5, 10, 20, 25 wt%) and ciprofloxacin (CIP: 1, 2.5, 5, 7.5, 20, 25), double-antibiotic (DA, MET+CIP) and triple-antibiotic (TA, MET+CIP+minocycline) series at nominal levels 5, 7, 9 and 11, Ca(OH)_2_ at 2–5, and a drug-free PVP control: 25 formulations with an outcome record. Twenty form fine fibers with reported means of 257–1873 nm. Five are unfavorable and are kept as data: MET 20 and 25 gave coarse microfibers (10.2 and 7.8 *µ*m), CIP 20 and Ca(OH)_2_ 5 gave no fibers, and CIP 25 gave sparse, enlarged fibers with no reported mean. Negative results carry the decision boundary for a formation classifier and are usually discarded; here they are retained. The full table is in Appendix A.

### What the audit established

(i) The DA 5% mean (804 *±* 255 nm, thesis p. 60) is absent from the published article and is restored, giving 20 diameters. (ii) CIP 25% gave sparse, enlarged fibers with no reported mean rather than no fibers at all; it is labeled unfavorable. (iii) The thesis gives a 0–40 kV supply range and room temperature rather than per-run settings, so we do not assert a fixed voltage, temperature or humidity. (iv) Three retained means exceed 1 *µ*m (PVP 1087, TA 5 1657, TA 7 1873 nm), so we call the binary target *fine-fiber formation* (mean below 2 *µ*m) rather than nanoscale spinnability; a strict sub-micron label is analyzed separately (17 positives among 24 classifiable rows; CIP 25 lacks the mean needed). (v) For DA and TA the thesis methods describe *each* antibiotic at the stated level, whereas figure labels use one percentage for the mixture. DA at level 9 may therefore mean 9% MET plus 9% CIP, not 9% total drug. We keep the nominal label and a per-component field and treat neither as a verified dry-mat drug fraction. We also correct a one-step label shift in the published diameter table, assign two transposed ATCC numbers as in the source’s Methods, and treat MIC values reported as *>* 1000 *µ*g mL^*−*1^ as right-censored.

### Biological summaries

Six groups have all three measured objectives: double and triple antibiotic pastes (DAP, TAP), the corresponding PVP fiber mats (DA-NFs, TA-NFs), and Ca(OH)_2_ as paste and as fibers. For each we take the worst-case minimum inhibitory concentration over four organisms (two *E. faecalis* strains, *E. coli, P. aeruginosa*), an ordinal 72 h MTT cytocompatibility score on NIH/3T3 cells encoded from the stated statistics (2: no significant reduction at either dose; 1: reduction at the high dose only; 0: reduction at both), and the absolute deviation of the four-week dentin phosphate/amide ratio from the untreated value of 10.446. These are single aggregate summaries without replicate-level data, and no viability percentage is invented anywhere in this work.

## 4 Stage 1: composition *→* morphology

### Models and evaluation

We use three encodings: **E1**, the nominal level plus a one-hot family indicator; **E2**, the four per-component concentrations plus family; and **E3**, composition descriptors (mass-weighted molecular weight, inorganic fraction, number of drugs, loading and log loading), with loading taken either as the nominal level or as the sum of components. Models are a majority predictor, a depth-one decision stump, L2 logistic regression (*C* = 1), a random forest (500 trees, minimum leaf size 2), ridge regression on log_10_ diameter (*α* = 1) and a Gaussian process (RBF plus white noise), all in scikit-learn [Pedregosa et al., 2011] with fixed settings and no tuning. We evaluate by leave-one-out (LOO) and by holding out whole formulation families, and report Wilson intervals, exact McNemar tests against the majority baseline, label-permutation tests (1000 permutations) and bootstrap intervals over out-of-fold predictions (5000 resamples). *R*^2^ is pooled over all out-of-fold predictions on the nanometer scale.

### Fine-fiber formation: what the classifier learned

Logistic regression classifies 24 of 25 formulations correctly (Table 1). So does a stump on the nominal level alone, and it keeps that accuracy when whole families are held out: every unfavorable outcome but one sits at level 20 or above, and every favorable one at 11 or below. The labels carry real signal (permutation *p* = 0.001), but the gain over the majority baseline is not established, because four discordant pairs against none give exact McNemar *p* = 0.125. Five failures are too few. The single error, Ca(OH)_2_ 5%, is a boundary inside a family whose training rows contain no failure. Ca(OH)_2_ is an inorganic particle that does not dissolve in ethanol, so its limit is not the organic-drug loading limit, and no model trained on the other 24 rows can see it.

**Table 1:** Morphology models. Top: fine-fiber formation (*n* = 25, 20 favorable). Bottom: mean diameter of the 20 retained formulations. Brackets are 95% intervals (Wilson for accuracy, bootstrap for MAE and *R*^2^). Encodings: E1 nominal level + family; E2 per-component concentrations + family; E3 composition descriptors with nominal (n.) or summed (s.) loading. “Family-out” holds out each formulation family in turn.

| Model | Input | LOO accuracy | Balanced | Family-out acc. |
| --- | --- | --- | --- | --- |
| Majority | – | 0.80 [0.61, 0.91] | 0.50 | 0.80 |
| Stump | nominal level | 0.96 [0.80, 0.99] | 0.90 | 0.96 |
| Stump | summed level | 0.60 [0.41, 0.77] | 0.38 | 0.60 |
| Logistic | E1 | 0.96 [0.80, 0.99] | 0.90 | 0.96 |
| Logistic | E2 | 0.84 [0.65, 0.94] | 0.60 | 0.72 |
| Logistic | E3 n. / s. | 0.96 / 0.88 | 0.90 / 0.70 | 0.92 / 0.84 |
| Random forest | E1 | 0.96 [0.80, 0.99] | 0.90 | 0.80 |

| Model | Input | LOO MAE (nm) | LOO MAPE | LOO $R^2$ | Family-out $R^2$ |
| --- | --- | --- | --- | --- | --- |
| Global geometric mean | – | 259 [136, 417] | 37.7% | –0.12 | –0.32 |
| Family geometric mean | – | 211 [106, 338] | 25.4% | 0.25 | –0.32 |
| Ridge | E1 | 193 [96, 317] | 23.4% | 0.34 [–0.35, 0.61] | –0.38 |
| Ridge | E3 n. | 193 [100, 309] | 22.6% | 0.38 [–0.31, 0.62] | –0.36 |
| Random forest | E1 | 205 [100, 332] | 25.5% | 0.27 [–0.94, 0.59] | –0.14 |
| Gaussian process | E1 | 159 [76, 260] | 18.5% | 0.53 [–0.16, 0.86] | –0.48 |

### The result depends on how a mixture is encoded

If the DA and TA levels mean per-component amounts, mixtures reach 22–33% total drug and still form fine fibers, while single drugs fail at 20%. A stump on summed loading then falls to 60%, component-encoded logistic regression to 84% (72% family-out), and summed descriptors to 88%. Which reading is correct is a recipe question the thesis leaves open (§3), and one extraction assay per mixture would settle it. Under a strict sub-micron target, logistic regression reaches 70.8% (17/24), no better than the majority. The 96% headline therefore predicts an operational campaign label at one loading interpretation. It is not a general measure of nanoscale spinnability.

### Diameter: interpolation, not transfer

Ridge regression reaches 193 nm MAE (95% CI 96–317) and *R*^2^ = 0.34 (*−* 0.35 to 0.61; permutation *p* = 0.002). That beats a global geometric mean (259 nm) and only modestly beats a per-family mean (211 nm). The Gaussian process has the best LOO error (159 nm, 18.5% MAPE, *R*^2^ = 0.53) and the worst family transfer (*R*^2^ = *−* 0.48). Every model has negative *R*^2^ when a family is held out, so the surrogates interpolate within the families they have seen and nothing more. Per-family LOO errors span an order of magnitude (ridge): 50 nm for DA, 72 for CIP, 86 for Ca(OH)_2_ and 129 for MET, against 566 for TA and 345 for the lone PVP control (Fig. 1b). The TA mats also have the widest fiber-to-fiber spread (SD 465–988 nm), so their mean is the least stable target. For a self-driving laboratory the useful output is this error map. The model can be trusted inside the MET, CIP, DA and Ca(OH)_2_ families at the sampled levels, and the TA family and anything outside the sampled families should be treated as unexplored.

**Figure 1:**
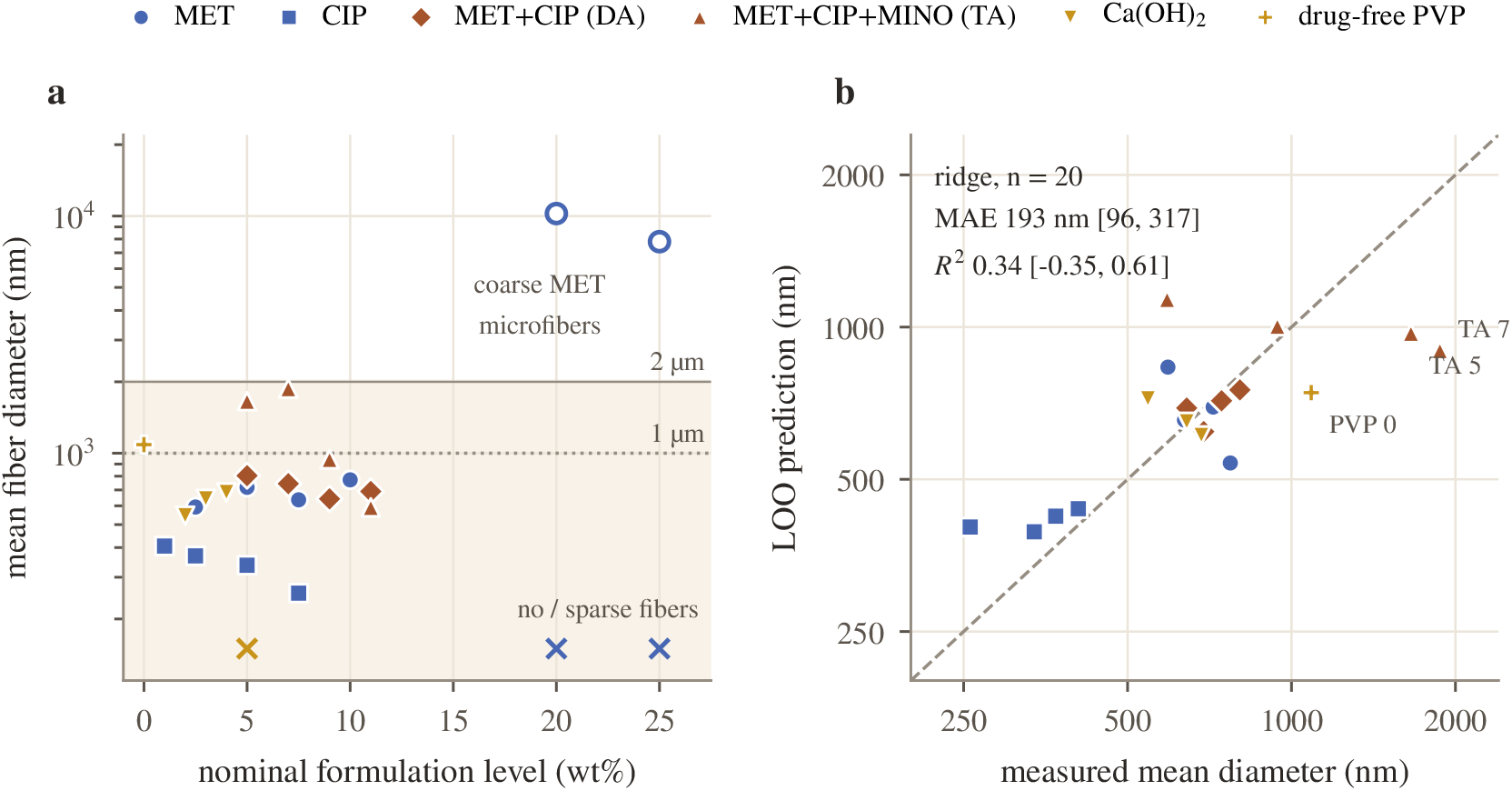
**(a)** The measured design space: filled markers are the 20 retained formulations, open markers the two coarse MET microfiber outcomes, crosses the formulations without a quantified diameter (plotted at the axis floor). The nominal level is a recipe label, not verified total drug loading. **(b)** LOO ridge predictions for the 20 retained formulations, with bootstrap 95% intervals. Errors are small where a family is densely sampled and large for the triple-antibiotic mats and the lone drug-free control.

### What the thesis-only runs bought

On a fixed test set of all 20 formulations, training on the 16 published diameters gives ridge MAE 200 nm and *R*^2^ = 0.30; adding the four thesis-only runs gives 193 nm and 0.34. The DA error falls from 68 to 50 nm, but the TA error barely moves (581 to 566 nm, and it rises from 573 to 602 nm for the random forest). Holding the test set fixed matters here: comparing models evaluated on different subsets would credit the recovered runs with gains that come from a changed target distribution. The recovered runs make the dataset more complete and only modestly more accurate. Adding the two MET microfiber means raises ridge MAE to 853 nm (53% MAPE), so the retained-group regression must not be read as a model of all morphologies.

## 5 Stage 2: same-drug evidence for a carrier-limited release risk

### Why *t*_90_

The campaign has no release curve, so whatever we transfer must be something the literature reports consistently. Published curves differ in time grid, medium and denominator, and are often shown only as figures. A time-to-fraction endpoint is a model-independent summary used to compare dissolution profiles [Costa and Sousa Lobo, 2001], and the time to 90% cumulative release, *t*_90_, asks the clinical question directly: has nearly all of the payload left the mat before the dressing interval ends? It can also be bounded from partial reports. If release was complete by time *T*, then *t*_90_ *≤ T* (left-censored); if less than 90% had been released at the last observation *T*, then *t*_90_ *> T* (right-censored), and we record the fraction reached. The cost is that *t*_90_ says nothing about local concentration, and it is undefined for a carrier that never releases 90%. Mechanistic models [Siepmann and Siepmann, 2012] need curve-level data that these reports do not share.

### Restricting to the same drugs

Pooling studies in which hydrophilic and hydrophobic carriers also carry different drugs confounds carrier with drug identity. We therefore built a table restricted to the two antibiotics in our mats, MET and CIP. This was a targeted search for electrospun fibers with quantitative *in vitro* release, not a systematic review. We included a report when a *t*_90_ bound could be read from its text or tables, without digitizing curves, and recorded medium, loading, the exact statement and the level of verification (Appendix B). Seventeen quantitative rows from seven studies qualify; three further studies contribute qualitative statements only.

### Results

The pattern in Fig. 2 and Table 2 has three parts. *Water-soluble carriers release fast*. All five PVP and PVA rows completed release within the observation window: PVP within *≤* 8 h, and in one study 94% within 5 min [Uhljar et al., 2021, Li et al., 2019]; thick PVA mats within about 80 h [Jannesari et al., 2011]. *Water-insoluble monolithic carriers release slowly and incompletely*. None of the eight ethyl cellulose, PVAc, PCL, PMMA or PLA rows reached 90% within windows of 3 h to 80 days, and the fraction released at the last report ranged from 1.5% to 50% [Li et al., 2019, Jannesari et al., 2011, Zupančič et al., 2016a, Reise et al., 2012, Adamczyk et al., 2025]. *Architecture fills the gap*. Blended fibers, core–shell fibers and a thicker mat released over 3 to 28 days, mostly to completion [Jannesari et al., 2011, Zupančič et al., 2016a,b]. For one PVA:PVAc blend, raising mat thickness from 100–140 to 680–760 *µ*m stretched total release from 72 to 325 h. Two studies compare carriers for the same drug at matched loading. In all four loading-matched pairs the insoluble carrier’s lower bound exceeds the soluble carrier’s upper bound: PVP *≤* 8 h versus ethyl cellulose *>* 72 h at 5% and 15% CIP [Li et al., 2019], and PVA *≤* 80 h versus PVAc *>* 1920 h at 5% and 10% CIP [Jannesari et al., 2011]. Two studies are too few for a significance test, but the direction is the same in both, and drug identity is held fixed by construction.

**Table 2:** Same-drug evidence by carrier group, and what it decides for a one-week *t*_90_ scenario. “Fails”: the bound places *t*_90_ under 168 h. “Exceeds”: *t*_90_ *>* 168 h, but release was incomplete. “Undecided”: the bound does not settle the question.

| Carrier group | Rows (studies) | $t_{90}$ bounds (h) | Released | One-week scenario | |
| --- | --- | --- | --- | --- | --- |
|  |  |  |  | fails / exceeds | undecided |
| Water-soluble (PVP, PVA) | 5 (3) | $\leq 0.08$ to $\leq 80$ | complete | 5 / 0 | 0 |
| Water-insoluble, monolithic | 8 (5) | $> 3$ to $> 1920$ | 1.5–50% | 0 / 5 | 3 |
| Blend, core-shell, thick mat | 4 (3) | $\leq 72$ to $\leq 672$ | complete | 1 / 0 | 3 |

**Figure 2:**
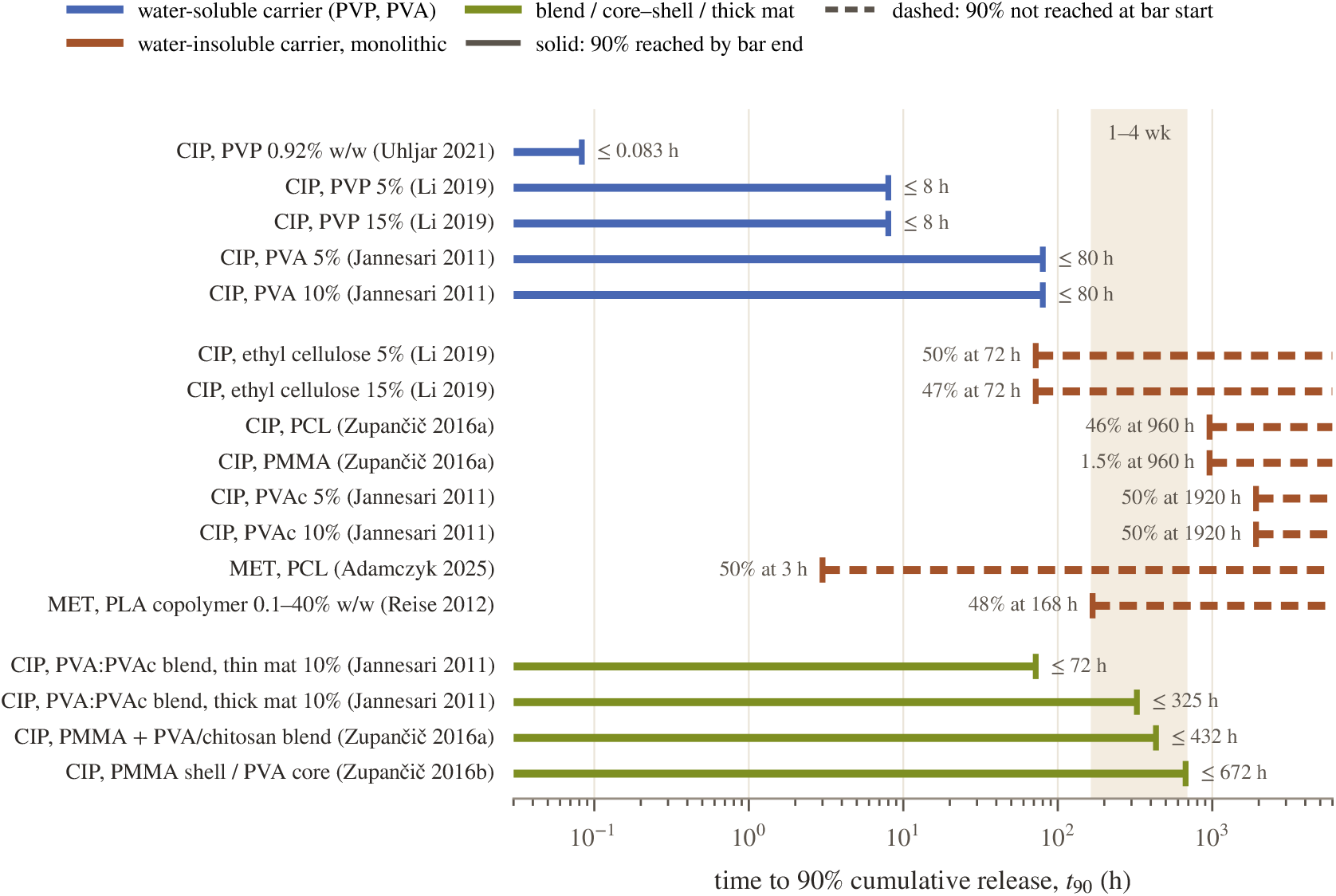
Same-drug release evidence for MET and CIP fibers. Solid bars: release was complete by the bar end (*t*_90_ *≤* bound). Dashed bars: 90% was not reached by the bar start (*t*_90_ *>* bound), labeled with the fraction released at that time. Shading marks a one-to-four-week scenario [American Association of Endodontists, 2022], which is a scheduling interval rather than a validated release requirement. Sources and assay conditions are listed in Table 6.

### What this supports, and what it does not

For the endodontic mats, which are all PVP, every same-drug PVP observation completed release within 8 h under sink conditions, at least 21 times shorter than a one-week scenario. Optimizing drug, loading or diameter within PVP is therefore unlikely to extend release to the dressing interval; the carrier or the fiber architecture has to change. We call this a *carrier-limited release risk* rather than a ceiling, for four reasons. First, slow carriers often release too little: PMMA released 1.5% and PCL 46% of CIP over 40 days [Zupančič et al., 2016a], which trades one failure (early depletion) for another (under-dosing). Second, insoluble does not mean slow: PCL released about half of its MET within 5 min at high loading [Adamczyk et al., 2025]. Third, geometry matters. In an outflow model of the eye, both PVP and PCL fibers extended CIP release [Baskakova et al., 2016], and a root canal is also a small, poorly exchanged volume, so a fast-dissolving mat may still sustain local exposure. Fourth, the one-to-four-week interval schedules a second visit [American Association of Endodontists, 2022]; it does not specify a release profile. The evidence also does not support the stronger claim that *no* formulation in this design space could reach the window. PVA mats 300–360 *µ*m thick came within about a factor of two of a week, and architecture closes the rest of the gap. What remains is a specific, falsifiable risk for PVP, and a single dissolution experiment can confirm or overturn it (§8).

## 6 Stage 3: a descriptive multi-objective screen

### Objectives and censoring

For the six groups with all three endpoints we maximize antimicrobial potency, *−*log_2_ (worst-case MIC), the ordinal cytocompatibility score, and dentin integrity, *−*| ΔP*/*A|. Six rounded summaries support a description, not an estimate, and we present the screen that way. Ca(OH)_2_ MICs are right-censored, so their potency is bounded above, not known. We call a group *necessarily* nondominated if no other group can dominate it for any MIC values allowed by the censoring, and *possibly* nondominated if it escapes domination for at least one allowed assignment.

### Partial order

DAP dominates TAP and DA-NFs dominates TA-NFs (Table 3, Fig. 3a). DAP, DA-NFs and Ca(OH)_2_ paste are necessarily nondominated. Ca(OH)_2_ fibers are possibly nondominated: they escape domination only if their true MIC is lower than the paste’s. Treating both censored values as 1000 would have forced a ranking that the data do not contain. In a related study, Ca(OH)_2_-loaded PVP fibers coated on gutta-percha were more active than the paste against *E. faecalis*, and neither inhibited *E. coli* or *P. aeruginosa* [Brimo et al., 2026], which is consistent with the censored worst-case MICs here. We treat one two-fold dilution as the plausible error of a single MIC readout. Across all 729 combinations of *±* 1 dilution on every MIC, the three necessary groups stay necessary in every scenario, Ca(OH)_2_ fibers stay possible in every one, TA-NFs is never nondominated, and TAP escapes domination only when its MIC shifts below DAP’s (33%). DA-NFs is the only fiber in the necessary set, which agrees with the source study’s expert reading.

**Table 3:** Six biological summaries from the source article [Brimo et al., 2023]. Cytocompatibility is ordinal (72 h MTT). Robustness: share of the 3^6^ = 729 combinations of a*±* 1 two-fold-dilution error on every MIC in which the group stays necessarily (possibly) nondominated.

| Group | Worst MIC ($\mu\text{g/mL}$) | Cyto | $\vert \Delta P/A\vert $ | Status | Robustness |
| --- | --- | --- | --- | --- | --- |
| DAP (paste) | 31.25 | 0 | 0.57 | necessary | 100% (100%) |
| DA-NFs (fiber) | 125 | 1 | 1.16 | necessary | 100% (100%) |
| TAP (paste) | 31.25 | 0 | 2.28 | dominated by DAP | 33% (33%) |
| TA-NFs (fiber) | 125 | 0 | 2.44 | dominated by DA-NFs | 0% (0%) |
| Ca(OH)<sub>2</sub> paste | > 1000 | 2 | 0.02 | necessary | 100% (100%) |
| Ca(OH)<sub>2</sub>-NFs (fiber) | > 1000 | 2 | 0.71 | possible | 0% (100%) |

**Figure 3:**
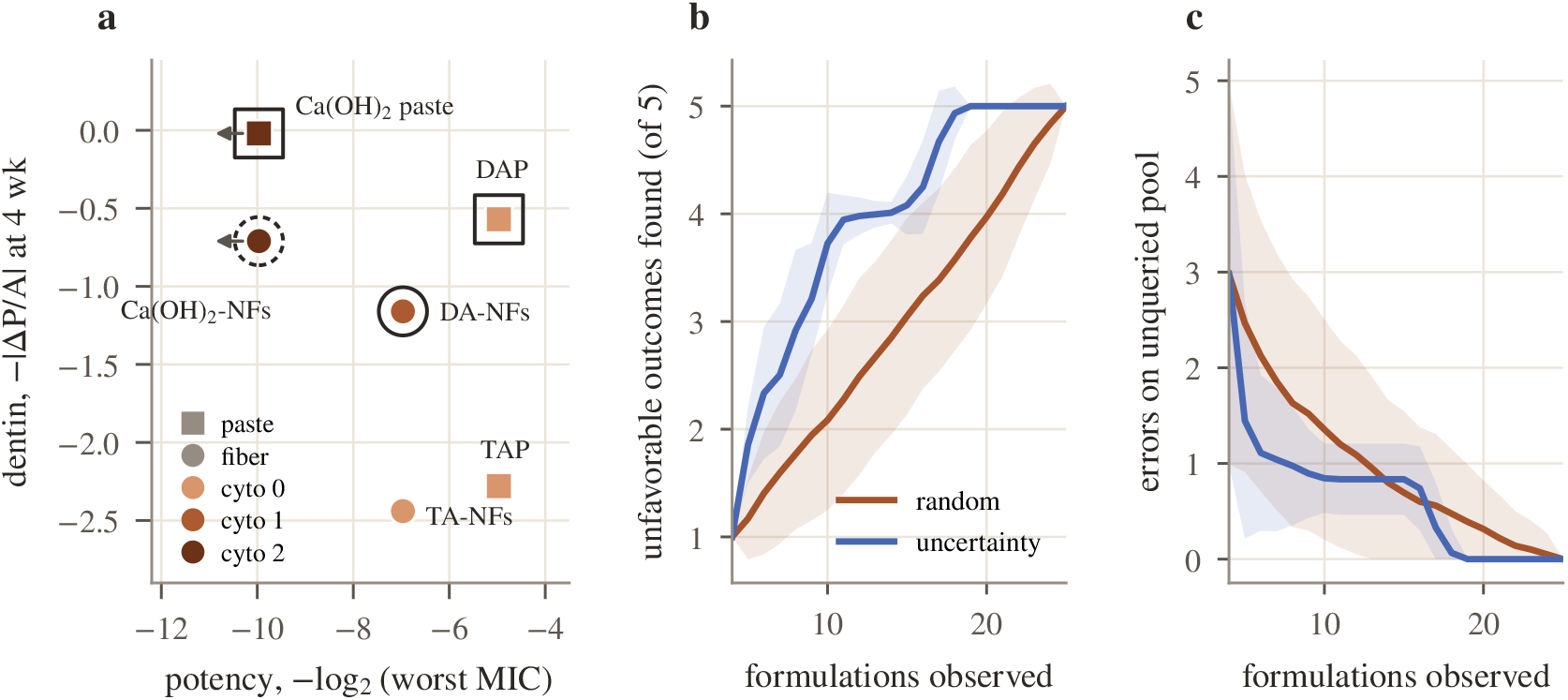
**(a)** Measured objectives for the six groups; squares are pastes, circles PVP fibers, fill encodes the ordinal cytocompatibility score, a solid outer ring marks necessarily and a dashed ring possibly nondominated groups, and arrows mark right-censored MICs. **(b, c)** Fixed-budget acquisition replays (200 paired seeds, mean *±* SD): unfavorable formulations discovered and errors on the unqueried pool.

### What the fiber format buys

Pastes and fiber mats have been compared qualitatively [Brimo et al., 2022]; the summaries here allow a coarse quantitative comparison. At matched drug system the fiber costs a factor of four in worst-case MIC for DA and TA (two dilutions, more than the one-dilution tolerance above). It also deviates more from untreated dentin in all three systems (by 0.59, 0.16 and 0.69). The cytocompatibility gain is specific to DA (score 1 versus 0); TA fibers reduce viability at both doses, as their paste does, which fits the field’s suspicion that minocycline is the problem. The fiber format therefore does not buy cytocompatibility in general. These are differences in encoded summaries, and paste and mat doses are defined differently, so none of them is a dose-normalized carrier effect.

### Release as a flag, not a filter

Imposing the transferred release constraint on every group would empty the front, but it would also apply a PVP release rule to paste vehicles, which contain no PVP. We therefore attach the Stage-2 risk only to the PVP fiber groups, as a flag. The result is specific: DA-NFs is the Pareto candidate, and its clinical viability depends on the one property nobody measured.

## 7 Stage 4: acquisition replays at fixed budgets

We replay the campaign as a sequential design problem. Each of 200 seeds starts from the same three favorable and one unfavorable formulation for both policies and refits the E1 logistic model after every query. Random acquisition follows a seeded order; uncertainty acquisition picks the pool formulation whose predicted probability is closest to 0.5. A tempting retrospective rule stops when the classifier becomes perfect on the unqueried pool. That rule reads labels an operating laboratory would not have, and under it the two policies look alike (median stops of 15 and 17 formulations).

At fixed budgets the picture reverses (Fig. 3b,c). After 10 formulations, uncertainty sampling has found 3.73 of the 5 unfavorable outcomes against 2.08 for random acquisition (paired difference 1.64, 95% CI 1.51–1.77). It has found all five by 20, when random acquisition has found 3.97. It also leaves fewer pool errors at most budgets; random acquisition is marginally better only around 14–16 formulations (difference 0.14 [0.03, 0.26] at 16). The plateau has a cause worth knowing. Uncertainty sampling finds the four high-loading failures early (median discovery at 5 to 10 formulations) but reaches Ca(OH)_2_ 5% only at a median of 17, later than random acquisition (14). The model assigns that formulation a 0.94 probability of success, so it is never uncertain about it, only wrong. An observable stopping rule, stopping after three consecutive queries whose labels the model had predicted correctly, halts at a median of 7 to 8 formulations, and in every replay it stops before all five failures are found. For small campaigns, uncertainty sampling maps the boundary the model already represents. Finding a boundary it does not represent takes coverage or chemical knowledge, and a stopping rule should be judged by what it misses.

## 8 Decision support: the experiment that decides

The analysis reduces the next campaign to one measurement on one formulation: release of MET and CIP from the DA-PVP mat, under sink conditions and in a confined volume that approximates a canal, alongside a same-drug comparator with a water-insoluble carrier or a barrier architecture. Table 4 lists what each outcome would mean before the experiment is run, so the result changes a decision rather than adding a data point. Appendix E gives the protocol, including loading verification by extraction, which also resolves the encoding question of §4.

**Table 4:** Pre-specified reading of the decisive experiment on the DA-PVP mat.

| Outcome | Reading | Next action |
| --- | --- | --- |
| $t_{90} \leq 24$ h (sink) and confined concentration below MIC before day 7 | risk confirmed | change carrier or architecture for the same drugs; stop tuning loading in PVP |
| $t_{90} \leq 24$ h (sink) but confined concentration ≥ MIC through day 7 | geometry sustains exposure | keep PVP; set dose for cytocompatibility |
| $24 < t_{90} \leq 168$ h | partly sustained | tune mat thickness or blend toward the target |
| $t_{90} > 168$ h with $\geq 90\%$ recovered by day 28 | risk refuted | proceed to eluate microbiology and cell assays |
| $< 50\%$ recovered by day 28 | availability problem | reformulate for complete release |

## 9 Limitations

This is a secondary analysis of one laboratory’s formulation summaries. There are no independent fabrication repeats, the standard deviations describe fibers within one mat, the recipe interpretation and tablet excipients are unresolved, and process settings were not recorded per run. Holding out families tests one kind of extrapolation; it does not test a new laboratory, solvent or polymer. The release table comes from a targeted search, not a systematic review. It has seven quantitative studies, two of which compare carriers within one study, with heterogeneous media, apparatus, mat thicknesses and loadings. Bounds were read from text and tables only: the numeric values for Li et al. [2019] come from its accepted manuscript, and the two Zupančič et al. [2016a,b] rows are approximate abstract-level bounds. The biological screen rests on six rounded summaries with an ordinal score that compresses dose and power, and the phosphate/amide ratio is a compositional proxy for dentin integrity. The acquisition replays reuse one finite dataset with oracle-stratified seeds. No dissolution, microbiological or cell experiment was performed for this paper.

## 10 Conclusion

A provenance-first reanalysis of a 25-formulation campaign sharpens both the dataset and what it can decide. The morphology classifier’s 96% is matched by a single threshold and depends on how mixtures are encoded; diameter surrogates interpolate within families and fail across them. Same-drug evidence for metronidazole and ciprofloxacin supports a carrier-limited release risk for PVP mats and identifies architecture, not loading, as the lever, but it does not support a hard ceiling. The screen narrows the candidates to one fiber, DA-NFs, whose viability hinges on one unmeasured property. For a field where each data point costs days of bench work, an analysis that names the next experiment and says in advance what each outcome would mean is worth more than a better interpolator.

## Data and code availability

The three data files (formulations with page-level provenance, biological summaries, and the release evidence table with exact statements and verification level) and the scripts that regenerate every number, table and figure in this paper are released with this paper. No replicate measurement is fabricated anywhere in this work.

## A EndoDesign-25: complete morphology table and provenance

All values are reproduced from the source thesis [Brimo, 2020]; page numbers are printed thesis pages. No value is read from a plotted bar height, and no replicate measurement is synthesized. Standard deviations describe fiber-to-fiber spread within one mat, not variation between independently fabricated mats, and are carried as metadata only.

**Table 5:** The 25 outcome-linked formulations. Level is the nominal recipe label in wt%; for DA and TA the thesis methods describe each antibiotic at that level. Pub.: diameter appeared in the published article [Brimo et al., 2023]. NR: not reported.

| Record | Level | Mean (nm) | SD (nm) | Outcome | Label | Pub. | Page |
| --- | --- | --- | --- | --- | --- | --- | --- |
| PVP_0 | 0 | 1087 | 285 | fibers | retained | yes | 48 |
| MET_2.5 / 5 / 7.5 / 10 | 2.5–10 | 593 / 718 / 636 / 772 | 105 / 148 / 121 / 192 | fibers | retained | yes | 50–51 |
| MET_20 | 20 | 10245 | 3480 | coarse microfibers | unfavorable | – | 45 |
| MET_25 | 25 | 7797 | 2670 | coarse microfibers | unfavorable | – | 45 |
| CIP_1 / 2.5 / 5 / 7.5 | 1–7.5 | 406 / 369 / 337 / 257 | 116 / 170 / 169 / 103 | fibers | retained | yes | 55–56 |
| CIP_20 | 20 | NR | NR | no fiber formation | unfavorable | – | 45–46 |
| CIP_25 | 25 | NR | NR | sparse, enlarged fibers | unfavorable | – | 46 |
| DA_5 | 5 | 804 | 255 | fibers | retained | no (recovered) | 60 |
| DA_7 / 9 | 7, 9 | 744 / 642 | 272 / 185 | fibers | retained | yes | 61 |
| DA_11 | 11 | 690 | 208 | fibers | retained | no | 61 |
| TA_5 | 5 | 1657 | 988 | fibers | retained | no | 65 |
| TA_7 / 9 | 7, 9 | 1873 / 942 | 985 / 682 | fibers | retained | yes | 65–66 |
| TA_11 | 11 | 591 | 465 | fibers | retained | no | 66 |
| CA_2 / 3 / 4 | 2–4 | 545 / 642 / 683 | 177 / 192 / 244 | fibers | retained | yes | 70 |
| CA_5 | 5 | NR | NR | no fibers | unfavorable | – | 70 |

**Table 6:**
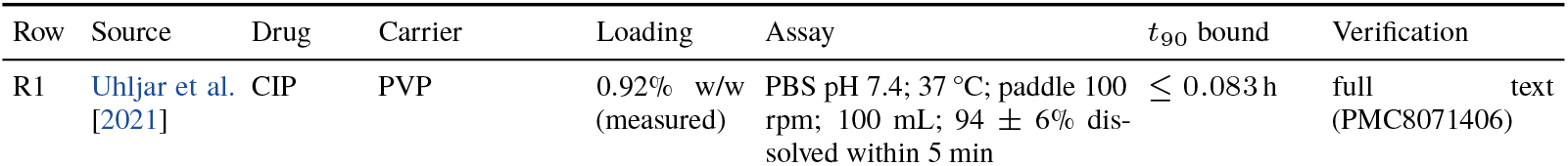
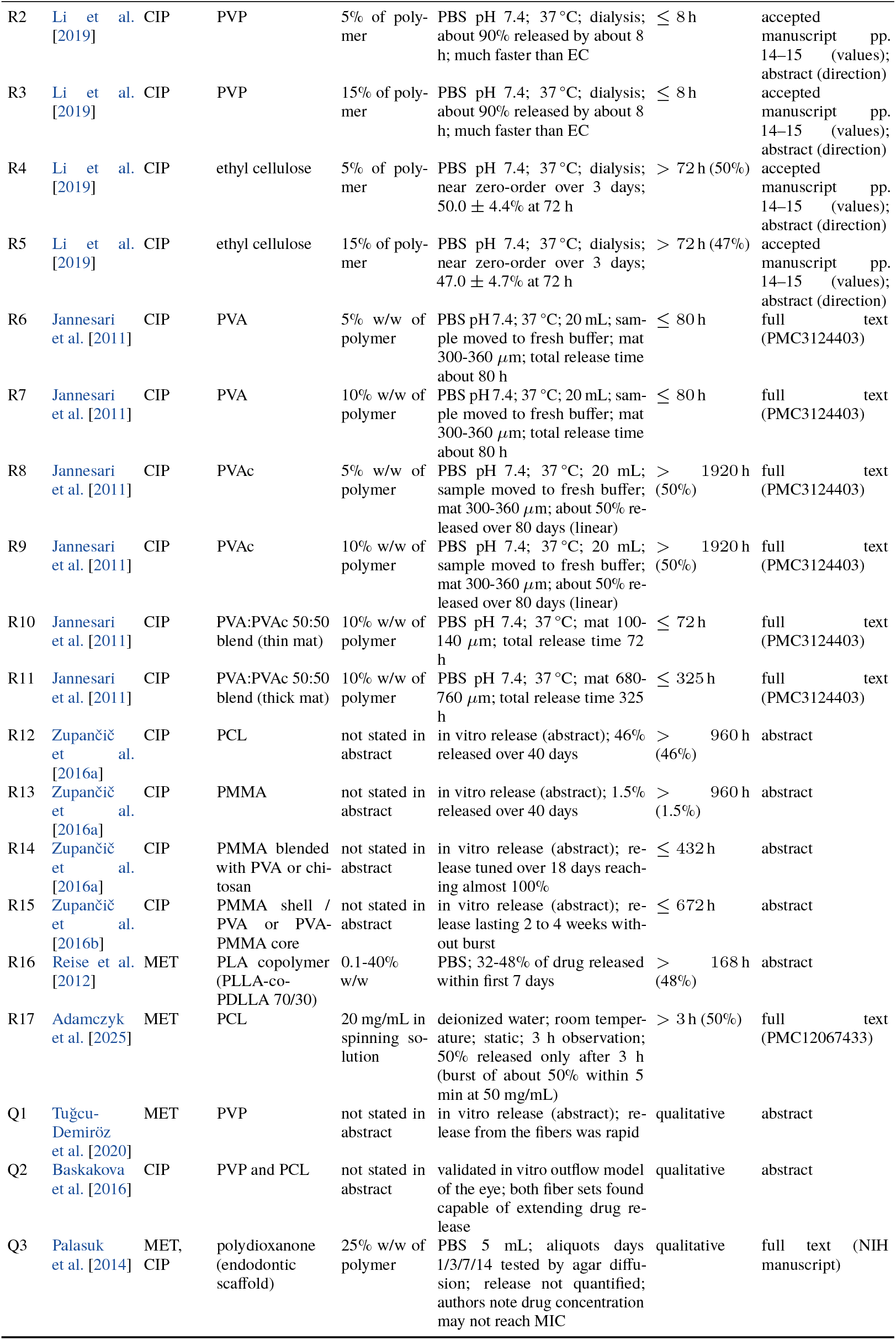
Same-drug release evidence (MET, CIP) used in §5.

### Provenance notes

(i) *Recovered diameter*. The DA 5% mean (804 *±* 255 nm) appears in thesis section 4.4.1 (p. 60) and is absent from the published article. (ii) *Label shift in the published article*. The article’s diameter table labels the DA and TA samples 9 and 11 wt% while its text and figure captions assign the same diameters to 7 and 9 wt%; the thesis runs both systems as complete 5/7/9/11 series and resolves the shift. (iii) *Loading definition*. The thesis methods describe each antibiotic in DA and TA at the stated level, while figure labels use one percentage for the mixture, and some solvent masses (DA 7%, TA 7%) do not reconcile cleanly with either reading. Overview prose mentions 10% where the detailed methods support 11%. We keep the nominal label and a per-component field side by side. Crushed pharmaceutical tablets were used, so excipients may contribute to the weighed powder. (iv) *Process metadata*. The thesis supports 8% PVP in ethanol, a nominal 0.9 mL h^*−*1^ flow rate, 15 cm working distance and room-temperature processing; it states a 0–40 kV supply range rather than a per-run voltage. A fixed voltage, temperature and humidity cannot be reconstructed per run and are not asserted. (v) *Unscorable preparations*. Preliminary Ca(OH)_2_ 20% and 25% recipes are mentioned (p. 30) without an outcome record; they are logged and excluded rather than counted as failures. (vi) *Biological summaries*. Worst-case MIC (four organisms), 72 h MTT categories and four-week dentin P/A deviations are aggregate summaries from the source article [Brimo et al., 2023]; the thesis lists biological testing as ongoing. Two ATCC numbers transposed in the source’s MIC table are assigned as in its Methods. MIC values reported as *>* 1000 *µ*g mL^*−*1^ are right-censored and never treated as 1000.

## B Same-drug release evidence: full table

Rows R1–R17 give a one-sided bound on *t*_90_; Q1–Q3 are qualitative. “Verification” states where the value was read. Numeric values for R2–R5 were extracted from the accepted manuscript of Li et al. [2019] (pp. 14–15); their direction (PVP much faster than ethyl cellulose; near zero-order ethyl cellulose release over three days) is confirmed by the published abstract. Abstract-level rows are approximate.

## C Additional morphology results

**Table 7:** All classification runs. McNemar *p*: exact two-sided test against the majority predictor. Errors list the misclassified formulations.

| Target | Model | Encoding | CV | Acc. | Bal. | $p$ | Errors |
| --- | --- | --- | --- | --- | --- | --- | --- |
| fine fiber | majority | E1 | LOO | 20/25 | 0.50 | 1.000 | MET 20, MET 25, CIP 20, CIP 25, CA 5 |
| fine fiber | majority | E1 | family | 20/25 | 0.50 | 1.000 | MET 20, MET 25, CIP 20, CIP 25, CA 5 |
| fine fiber | stump | level only | LOO | 24/25 | 0.90 | 0.125 | CA 5 |
| fine fiber | stump | level only | family | 24/25 | 0.90 | 0.125 | CA 5 |
| fine fiber | stump | summed only | LOO | 15/25 | 0.38 | 0.062 | MET 20, MET 25, CIP 20, CIP 25, DA 9, DA 11, TA 7, TA 9, TA 11, CA 5 |
| fine fiber | stump | summed only | family | 15/25 | 0.38 | 0.062 | MET 20, MET 25, CIP 20, CIP 25, DA 9, DA 11, TA 7, TA 9, TA 11, CA 5 |
| fine fiber | logistic | E1 | LOO | 24/25 | 0.90 | 0.125 | CA 5 |
| fine fiber | logistic | E1 | family | 24/25 | 0.90 | 0.125 | CA 5 |
| fine fiber | forest | E1 | LOO | 24/25 | 0.90 | 0.125 | CA 5 |
| fine fiber | forest | E1 | family | 20/25 | 0.50 | 1.000 | MET 20, MET 25, CIP 20, CIP 25, CA 5 |
| fine fiber | logistic | E2 | LOO | 21/25 | 0.60 | 1.000 | MET 20, MET 25, CIP 20, CA 5 |
| fine fiber | logistic | E2 | family | 18/25 | 0.45 | 0.500 | MET 20, MET 25, CIP 20, CIP 25, DA 11, TA 11, CA 5 |
| fine fiber | forest | E2 | LOO | 20/25 | 0.50 | 1.000 | MET 20, MET 25, CIP 20, CIP 25, CA 5 |
| fine fiber | forest | E2 | family | 20/25 | 0.50 | 1.000 | MET 20, MET 25, CIP 20, CIP 25, CA 5 |
| fine fiber | logistic | E3 nominal | LOO | 24/25 | 0.90 | 0.125 | CA 5 |
| fine fiber | logistic | E3 nominal | family | 23/25 | 0.80 | 0.250 | CIP 20, CA 5 |
| fine fiber | logistic | E3 summed | LOO | 22/25 | 0.70 | 0.500 | MET 20, CIP 20, CA 5 |
| fine fiber | logistic | E3 summed | family | 21/25 | 0.60 | 1.000 | MET 20, CIP 20, CIP 25, CA 5 |
| sub-micron | majority | E1 | LOO | 17/24 | 0.50 | 1.000 | PVP 0, MET 20, MET 25, CIP 20, TA 5, TA 7, CA 5 |
| sub-micron | majority | E1 | family | 17/24 | 0.50 | 1.000 | PVP 0, MET 20, MET 25, CIP 20, TA 5, TA 7, CA 5 |
| sub-micron | stump | level only | LOO | 20/24 | 0.71 | 0.250 | PVP 0, TA 5, TA 7, CA 5 |
| sub-micron | stump | level only | family | 18/24 | 0.57 | 1.000 | PVP 0, MET 20, MET 25, TA 5, TA 7, CA 5 |
| sub-micron | stump | summed only | LOO | 16/24 | 0.64 | 1.000 | PVP 0, DA 7, DA 9, DA 11, TA 5, TA 9, TA 11, CA 5 |
| sub-micron | stump | summed only | family | 12/24 | 0.39 | 0.125 | PVP 0, MET 20, MET 25, CIP 1, CIP 20, DA 7, DA 9, DA 11, TA 5, TA 9, TA 11, CA 5 |
| sub-micron | logistic | E1 | LOO | 17/24 | 0.58 | 1.000 | PVP 0, CIP 20, TA 5, TA 7, TA 9, TA 11, CA 5 |
| sub-micron | logistic | E1 | family | 20/24 | 0.71 | 0.250 | PVP 0, TA 5, TA 7, CA 5 |
| sub-micron | forest | E1 | LOO | 17/24 | 0.50 | 1.000 | PVP 0, MET 20, MET 25, CIP 20, TA 5, TA 7, CA 5 |
| sub-micron | forest | E1 | family | 17/24 | 0.50 | 1.000 | PVP 0, MET 20, MET 25, CIP 20, TA 5, TA 7, CA 5 |
| sub-micron | logistic | E2 | LOO | 15/24 | 0.44 | 0.500 | PVP 0, MET 20, MET 25, CIP 20, TA 5, TA 7, TA 9, TA 11, CA 5 |
| sub-micron | logistic | E2 | family | 14/24 | 0.41 | 0.250 | PVP 0, MET 20, MET 25, CIP 20, DA 7, DA 9, DA 11, TA 5, TA 7, CA 5 |
| sub-micron | forest | E2 | LOO | 17/24 | 0.50 | 1.000 | PVP 0, MET 20, MET 25, CIP 20, TA 5, TA 7, CA 5 |
| sub-micron | forest | E2 | family | 17/24 | 0.50 | 1.000 | PVP 0, MET 20, MET 25, CIP 20, TA 5, TA 7, CA 5 |
| sub-micron | logistic | E3 nominal | LOO | 19/24 | 0.64 | 0.500 | PVP 0, CIP 20, TA 5, TA 7, CA 5 |
| sub-micron | logistic | E3 nominal | family | 19/24 | 0.64 | 0.500 | PVP 0, CIP 20, TA 5, TA 7, CA 5 |
| sub-micron | logistic | E3 summed | LOO | 17/24 | 0.54 | 1.000 | PVP 0, MET 20, CIP 20, TA 5, TA 7, TA 11, CA 5 |
| sub-micron | logistic | E3 summed | family | 16/24 | 0.47 | 1.000 | PVP 0, MET 20, MET 25, CIP 20, DA 11, TA 5, TA 7, CA 5 |

**Table 8:**
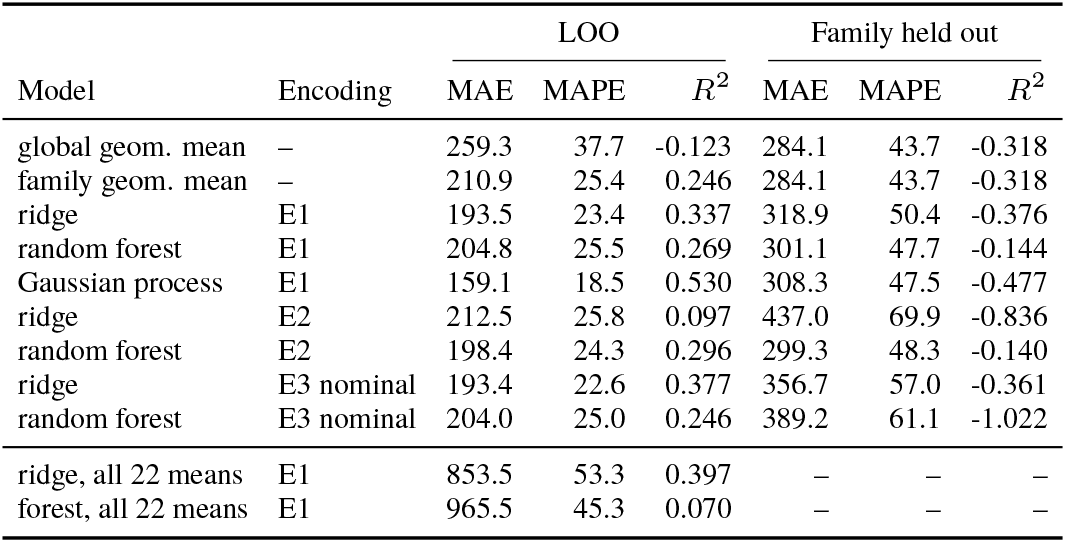
All diameter-regression runs (*n* = 20 retained formulations unless noted). MAE in nm, MAPE in %, *R*^2^ on the nanometer scale.

| Model | Encoding | LOO |  |  | Family held out |  |  |
| --- | --- | --- | --- | --- | --- | --- | --- |
| | | MAE | MAPE | $R^2$ | MAE | MAPE | $R^2$ |
| global geom. mean | — | 259.3 | 37.7 | -0.123 | 284.1 | 43.7 | -0.318 |
| family geom. mean | — | 210.9 | 25.4 | 0.246 | 284.1 | 43.7 | -0.318 |
| ridge | E1 | 193.5 | 23.4 | 0.337 | 318.9 | 50.4 | -0.376 |
| random forest | E1 | 204.8 | 25.5 | 0.269 | 301.1 | 47.7 | -0.144 |
| Gaussian process | E1 | 159.1 | 18.5 | 0.530 | 308.3 | 47.5 | -0.477 |
| ridge | E2 | 212.5 | 25.8 | 0.097 | 437.0 | 69.9 | -0.836 |
| random forest | E2 | 198.4 | 24.3 | 0.296 | 299.3 | 48.3 | -0.140 |
| ridge | E3 nominal | 193.4 | 22.6 | 0.377 | 356.7 | 57.0 | -0.361 |
| random forest | E3 nominal | 204.0 | 25.0 | 0.246 | 389.2 | 61.1 | -1.022 |
| ridge, all 22 means | E1 | 853.5 | 53.3 | 0.397 | — | — | — |
| forest, all 22 means | E1 | 965.5 | 45.3 | 0.070 | — | — | — |

**Table 9:** Value of thesis-only data on fixed test sets (E1 encoding). Top: all 20 formulations evaluated by LOO, training on the published 16 (minus the held-out row) or on all 20. Bottom: the 19 formulations other than DA 5%, with and without DA 5% in training.

| Model | Training set | MAE | MAPE | $R^2$ | DA MAE | TA MAE |
| --- | --- | --- | --- | --- | --- | --- |
| ridge | published 16 | 200.3 | 24.0 | 0.296 | 68.0 | 581.3 |
| ridge | all 20 | 193.5 | 23.4 | 0.337 | 50.0 | 566.1 |
| random forest | published 16 | 206.2 | 25.1 | 0.248 | 99.8 | 572.9 |
| random forest | all 20 | 204.8 | 25.5 | 0.269 | 48.5 | 601.5 |
| ridge | 19, without DA 5% | 203.3 | 24.6 | 0.326 | – | – |
| ridge | 19, with DA 5% | 200.9 | 24.3 | 0.337 | – | – |
| random forest | 19, without DA 5% | 209.6 | 25.9 | 0.257 | – | – |
| random forest | 19, with DA 5% | 213.1 | 26.5 | 0.269 | – | – |

## D Acquisition replays: details

**Table 10:** Fixed-budget replays, 200 paired seeds (mean *±* SD). Δ: uncertainty minus random, paired, with a 95% bootstrap interval over seeds.

| Observed | Unfavorable outcomes found (of 5) |  |  | Errors on unqueried pool |  |  |
| --- | --- | --- | --- | --- | --- | --- |
| | random | uncertainty | $\Delta$ | random | uncertainty | $\Delta$ |
| 6 | 1.41 ± 0.57 | 2.33 ± 0.61 | +0.93 [+0.82, +1.04] | 2.12 ± 1.42 | 1.11 ± 0.81 | -1.01 [-1.25, -0.80] |
| 8 | 1.77 ± 0.70 | 2.92 ± 0.75 | +1.15 [+1.01, +1.29] | 1.63 ± 1.29 | 0.97 ± 0.62 | -0.66 [-0.85, -0.47] |
| 10 | 2.08 ± 0.84 | 3.73 ± 0.47 | +1.64 [+1.51, +1.77] | 1.35 ± 1.15 | 0.84 ± 0.36 | -0.51 [-0.68, -0.35] |
| 12 | 2.49 ± 0.91 | 3.98 ± 0.17 | +1.49 [+1.36, +1.62] | 1.09 ± 1.04 | 0.83 ± 0.37 | -0.26 [-0.41, -0.10] |
| 14 | 2.85 ± 0.90 | 4.01 ± 0.10 | +1.16 [+1.03, +1.28] | 0.80 ± 0.86 | 0.83 ± 0.37 | +0.04 [-0.09, +0.15] |
| 16 | 3.24 ± 0.86 | 4.25 ± 0.43 | +1.01 [+0.89, +1.14] | 0.60 ± 0.78 | 0.74 ± 0.44 | +0.14 [+0.03, +0.26] |
| 18 | 3.58 ± 0.85 | 4.93 ± 0.25 | +1.36 [+1.24, +1.49] | 0.47 ± 0.67 | 0.07 ± 0.25 | -0.41 [-0.52, -0.31] |
| 20 | 3.97 ± 0.80 | 5.00 ± 0.00 | +1.03 [+0.92, +1.14] | 0.32 ± 0.51 | 0.00 ± 0.00 | -0.32 [-0.39, -0.24] |
| 22 | 4.43 ± 0.65 | 5.00 ± 0.00 | +0.57 [+0.47, +0.67] | 0.14 ± 0.36 | 0.00 ± 0.00 | -0.14 [-0.20, -0.09] |
| 24 | 4.83 ± 0.37 | 5.00 ± 0.00 | +0.17 [+0.12, +0.21] | 0.05 ± 0.22 | 0.00 ± 0.00 | -0.05 [-0.08, -0.03] |

### Discovery order and stopping

Median budget at which each unfavorable formulation is queried (random / uncertainty; the four seed formulations count toward the budget): MET 20 11 / 6; MET 25 13 / 8; CIP 20 12.5 / 5; CIP 25 12 / 10; CA 5 14 / 17. At a budget of 12, the model’s mean predicted probability that Ca(OH)_2_ 5% forms fine fibers is 0.94 (uncertainty runs) and 0.95 (random runs). An oracle rule (stop when the pool is predicted perfectly) gives median stops of 15 and 17 (means 14.8 and 15.6). The observable three-in-a-row rule stops at a median of 7 (random) and 8 (uncertainty); the share of runs that stop with at least one failure undiscovered is 100% and 100%, missing on average 3.29 and 2.15 of the five.

## E Proposed validation protocol (not performed here)

No dissolution, microbiological or cell-culture experiment was performed for this paper. The protocol below is the measurement that would decide Table 4.

### Materials and controls

The DA-PVP mat at the nominal level selected by the screen; drug-free PVP mat (blank); an equivalent free-drug MET+CIP mixture (dissolution control); and a same-drug comparator built on a water-insoluble carrier or a barrier architecture (core–shell or blended fiber, or a thicker mat). Match measured drug mass, antibiotic ratio, exposed area and mat mass where feasible and record the differences that remain. Use at least three independently fabricated mats per condition for a feasibility study, and set the final sample size from the observed between-batch variation.

### Analytics

Quantify MET and CIP separately with a validated chromatographic method (calibration range, limits of quantification, stability in medium, adsorption to vessels and membranes). Determine the actual loading of every batch by independent extraction; this also settles the nominal-versus-per-component ambiguity of §3.

### Release arms

(1) A buffered sink-condition assay at 37 °C, verifying sink conditions for both drugs rather than assuming them. (2) A confined-volume arm whose liquid volume and exchange rate approximate a root canal, so that local concentration, not only fractional release, is measured. Sample at 5, 15 and 30 min, 1, 2, 4 and 8 h, 1 and 3 d, and 7, 14 and 28 d; correct cumulative mass for withdrawn volume.

### Reporting

For each drug: recovered mass and mass balance (released plus residual extracted from the mat at the end), the fraction released at each scheduled time, concentration-time profiles in the confined arm, and *t*_50_ and *t*_90_ only when observed, otherwise the censoring time and the fraction reached. Test eluates against the four organisms of the source study and against relevant dental pulp or apical papilla cells under a prespecified exposure design.

